# Light-exercise-induced dopaminergic and noradrenergic stimulation in the dorsal hippocampus: using a rat physiological exercise model

**DOI:** 10.1101/2023.06.19.545490

**Authors:** Taichi Hiraga, Toshiaki Hata, Shingo Soya, Ryo Shimoda, Kanako Takahashi, Mariko Soya, Koshiro Inoue, Joshua P. Johansen, Masahiro Okamoto, Hideaki Soya

## Abstract

Exercise activates the dorsal hippocampus, which triggers the synaptic and cellar plasticity and ultimately promotes memory formation. For decades, these benefits have been explored using demanding and stress-response-inducing exercise at moderate-to-vigorous intensities. In contrast, our translational research with animals and humans has focused on light exercise below the lactate threshold (LT), which almost anyone can safely perform with minimal stress, and found that even light exercise can stimulate hippocampal activity and enhance memory performance. Although the circuit mechanism of this boost remains unclear, arousal promotion even with light exercise implies the involvement of the ascending monoaminergic system, which is essential to modulate hippocampal activity and impact memory. To examine this hypothesis, we employed our physiological exercise model based on the LT of rats that can be applied to human and immunohistochemically assessed the neuronal activation of the dorsal hippocampal sub-regions and brainstem monoaminergic neurons. Also, we monitored the dynamics of monoamine release at the dorsal hippocampus using *in vivo* microdialysis. We found that even light exercise increased neuronal activity in the dorsal hippocampal sub-regions and induced noradrenaline and dopamine release. Furthermore, we found that tyrosine hydroxylase-positive neurons in the locus coeruleus (LC) and the ventral tegmental area (VTA) were activated even by light exercise and were both positively correlated with the dorsal hippocampal activation. In conclusion, our findings demonstrate that light exercise stimulates hippocampal neurons, possibly through the LC-noradrenergic and/or VTA-dopaminergic neurons. This sheds light on the circuit mechanisms responsible for hippocampal neural activation during exercise, consequently enhancing memory function.

**Graphical abstract:** Our previous research with animals and humans has demonstrated that even light exercise can boost neuronal activity in the dorsal hippocampus and improve memory. While the mechanism underlying this remains undetermined, recent studies suggest the involvement of the ascending monoaminergic system. Here, we examined this hypothesis and found a possible contribution of noradrenergic neurons in the locus coeruleus and dopaminergic neurons in the ventral tegmental area to dorsal hippocampal activation during light exercise, implying a circuit mechanism for light-exercise-enhanced memory.

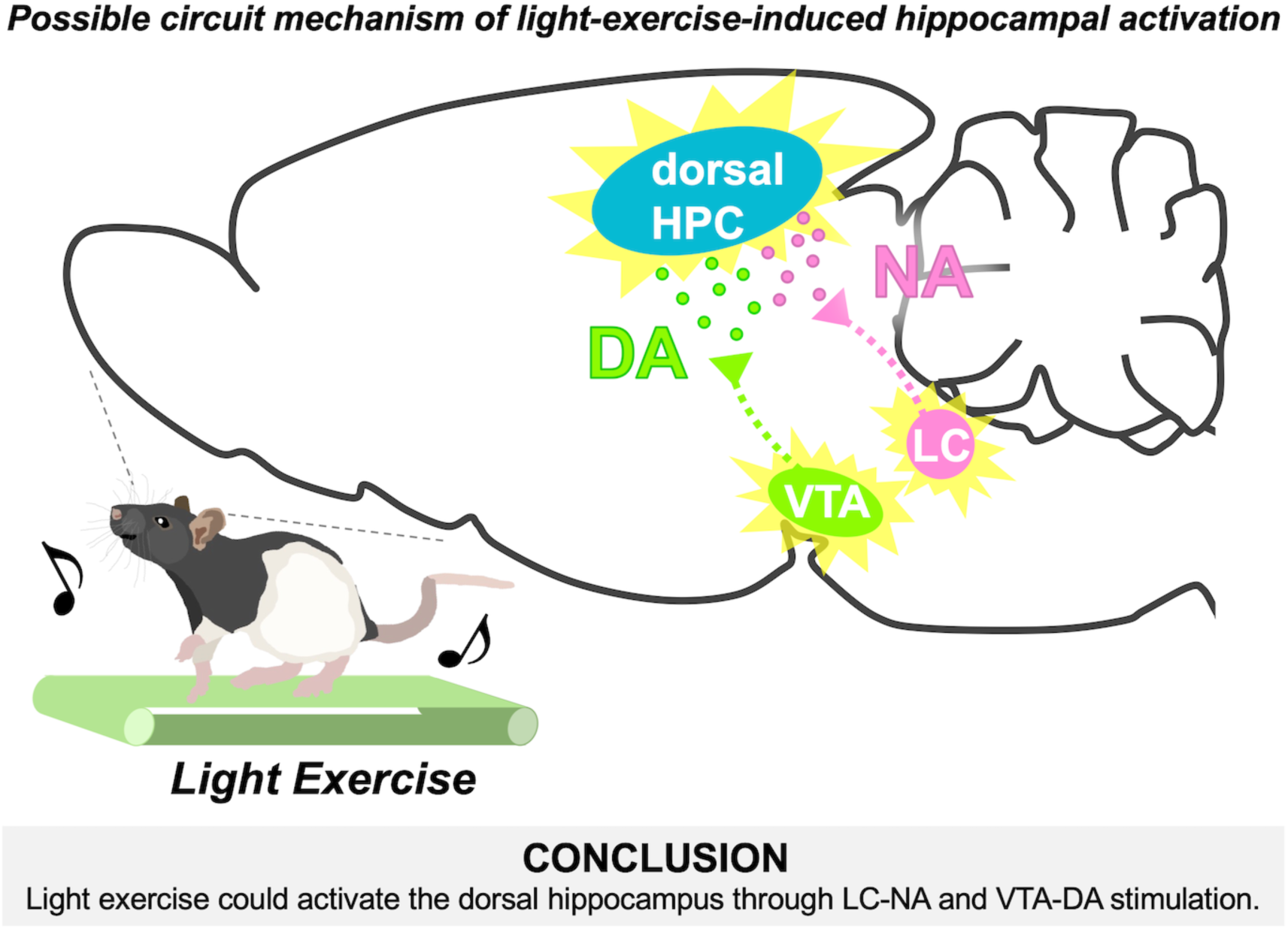

## 1 Introduction

There is accumulating evidence showing that physical exercise has beneficial effects on the brain and its functions, including hippocampus-dependent learning and memory.^1–4^ The hippocampus is well-known for being involved in several processes related to emotion, stress resilience, and cognitive function. In particular, the dorsal part of the hippocampus plays a key role in the learning process and memory formation.^5^ Physical exercise has been found to stimulate the activity of the dorsal hippocampal neurons.^6–8^ This subsequently triggers the production of plasticity-related proteins (PRPs) such as the brain-derived neurotrophic factor (BDNF) in the hippocampus.^9–11^ Also, this activation increases cerebral blood flow^8,12^ and facilitates the entrance of bioactive hormones “exerkines” secreted from peripheral organs, such as the insulin-like growth factor I (IGF-I), cathepsin B, and irisin, across the blood-brain barrier into the hippocampus.^13–18^ In turn, these boosts ultimately contribute to cellar and synaptic plasticity and memory enhancement.^1,15,16,19–21^

For decades, the benefits of exercise to learning and memory have been based on evidence using exercise at moderate-to-vigorous intensities,^2–4,22^ which is generally recommended in exercise therapy. However, such demanding exercise can cause physiological stress and reduce exercise adherence.^23–27^ Besides, while the chronic intervention of voluntary wheel running is widely used in animal research,^1,15,16,18^ this exercise model provides less information about effective exercise intensity, frequency, and duration for memory enhancement. Moreover, voluntary running rodents run at a relatively high intensity (45 m/min) and exhibit intermittent running patterns in the dark cycle,^28^ suggesting that wheel running is not an appropriate translational exercise model for human exercise. We thus have been employing a rodent treadmill running model based on physiological responses such as lactate threshold (LT). LT is the signpost for exercise-induced stress, which begins with the onset of lactate accumulation in the blood and the accumulation of adrenocorticotropic hormone (ACTH).^23,25^ This physiological exercise model enables us to define running below the LT as light exercise with minimum physiological stress and running above the LT as vigorous, stress-inducing exercise stimuli. Using this model, we found that even light exercise was sufficient to acutely stimulate hippocampal activity and BDNF induction,^9^ and to promote hippocampal neurogenesis and spatial memory.^29–31^ Conversely, excessive ACTH secretion induced by vigorous exercise cancels out these beneficial effects on hippocampal plasticity and memory due to the stress-induced hippocampal vulnerability.^29–33^ Additionally, our research with human participants and functional magnetic resonance imaging (fMRI) found that acute light exercise increases hippocampal activity and enhances pattern separation, which is a hippocampus-related function.^34^ Taken together, our translational research demonstrates the importance of exercise intensity for determining the effect of exercise on the hippocampus and highlights the advantages of light exercise.

However, prior studies, including our own findings, have yet to address the circuit mechanism underlying dorsal hippocampal activation by light exercise and its beneficial effect on memory performance. Classically, it is known that exercise can promote arousal,^35–37^ which is thought to originate from the activation of the ascending arousal system (AAS).^38,39^ This is supported by our recent studies showing that arousal promotion can occur by exercise even at very light-intensity accompanied by pupil dilation,^34,40–43^ which reflects the activity of AAS.^44–48^ The AAS sends out widespread projections in order to increase global brain activity, including the dorsal hippocampus, which largely overlaps the monoaminergic neuromodulatory system. Monoamines like noradrenaline (NA), dopamine (DA), and serotonin (5-HT) are essential for multiple functions such as arousal, attention, reward, motivation, and mood regulation.^49–55^ At the dorsal hippocampus, monoaminergic stimulation facilitates learning and memory processes by regulating the expression of PRPs and synaptic and cellar plasticity, such as long-term potentiation (LTP) and neurogenesis.^56–61^ The hippocampal monoamines are mainly synthesized and provided by the specific nuclei located in the pons noradrenergic locus coeruleus (LC),^62–66^ the midbrain dopaminergic nuclei, such as the ventral tegmental area (VTA) and the substantia nigra pars compacta (SNpc),^67–71^ and the midbrain serotonergic raphe nuclei (the dorsal raphe: DRN and the median raphe: MRN).^72,73^ Previous research demonstrate that single bouts of exercise activate these brainstem monoaminergic neurons^74–76^ and affect the hippocampal monoamine transmissions,^3,77,78^ thereby contributing to the enhancement of hippocampus-related functions.^3,74,79^ These findings lead us to the hypothesis that the monoamines provided by the brainstem nuclei could be involved in the light exercise-induced dorsal hippocampal activation, as the neural basis of memory enhancement.

To test this hypothesis, the involvement of the brainstem monoaminergic system in dorsal hippocampal activation during light exercise was investigated using a rat LT-based treadmill running model that can be applied to humans. We analyzed the neuronal activation of the dorsal hippocampus and of the brainstem monoaminergic neurons using immunohistochemistry. Also, we monitored the dorsal hippocampal monoamine transmission with *in vivo* microdialysis combined with the high-performance liquid chromatography (HPLC) system.

## 2 Materials and Methods

### 2.1 Animals

All animal care and experimental procedures were performed in accordance with protocols approved by the University of Tsukuba Animal Experiment Committee (19-378, 20-405, and 21-373) based on the NIH Guidelines for the Care and Use of Laboratory Animals (NIH publication, 1996). Male adult Long-Evans rats (*n* = 72; 380-470 g; 10-12 weeks old at the beginning of experiments) from the TH-Cre rat colony (i.e., Cre-negative littermate control of TH-Cre Long-Evans rats) were housed with 2–3 rats in each cage and singly housed after surgery. Rats were kept in a controlled environment at 22 ± 1°C with a 12/12-h light/dark cycle (lights on 7 a.m. to 7 p.m.) and given *ad libitum* access to water and food (MF, Oriental Yeast Co., Ltd., Tokyo, Japan). All experiments were done during the light cycle.

### 2.2 Experimental design

Four experiments were performed independently: measurement of LT (Experiment 1: *n* = 6), validation of exercise intensity (Experiment 2: *n* = 5-7/group, 3 groups), immunohistochemical analysis of neuronal activation in dorsal hippocampus and brainstem monoaminergic nuclei (Experiment 3: *n* = 6-8/group, 3 groups), and analysis of dorsal hippocampal monoamine release with *in vivo* microdialysis (Experiment 4: *n* = 6-7/group, 3 groups). All rats were handled for 1 week before the start of all procedures and habituated to treadmill running (6 days, Table 1). Then, they were randomly allocated to three groups based on LT as follows: 1) sedentary on treadmill (Control: 0 m/min), 2) below-LT light-intensity (low speed) running (Light: 15 m/min), and 3) above-LT vigorous-intensity (high speed) running (Vigorous: 25 m/min). Exercise and control conditions were maintained for 30 minutes.

**Table 1.** Protocol for habituation to treadmill running.

| Day | Running speed (duration) |
| --- | --- |
| 1 | Rest (10 min) → 5 m/min (10 min) → 10 m/min (10 min) |
| 2 | Rest (5 min) → 5 m/min (10 min) → 10 m/min (10 min) → 15 m/min (10 min) |
| 3 | Rest |
| 4 | Rest (5 min) → 10 m/min (10 min) → 15 m/min (10 min) → 20 m/min (10 min) |
| 5 | Rest (5 min) → 15 m/min (10 min) → 20 m/min (10 min) → 25 m/min (10 min) |
| 6 | Rest (5 min) → 15 m/min (10 min) → 20 m/min (10 min) → 25 m/min (10 min) |

### 2.3 Surgery to insert jugular catheter

In Experiments 1 and 2, 1 day after the habituation to treadmill running periods, the rats were anesthetized with isoflurane, and a silicone catheter was inserted into the jugular vein and fixed with a silk thread suture (37 mm). The external distal end of the catheter was fixed at the nape of the neck. After the surgery, the rats were injected with antibiotics (Mycillin Sol; Meiji Seika, Tokyo, Japan). Then, they were housed individually and were allowed to recover for 2 or 3 days before the treadmill running test.

### 2.4 Stereotaxic surgery

In Experiment 4, after a week of handling, all rats were anesthetized with isoflurane and placed in a stereotaxic frame (Model 51900, Muromachi Kikai Co., Ltd., Tokyo, Japan). A guide cannula (AG-6, Eicom, Kyoto, Japan) was implanted above the dorsal hippocampus (AP: -3.6 mm, ML: 3.8 mm, DV: -1.8 mm, 20° angle, from bregma). The cannula was secured to the skull with three anchoring screws and dental cement. To prevent occlusion, a dummy cannula (AD-6, Eicom) was inserted into the guide cannula. After the surgery, the rats were injected with antibiotics (Mycillin Sol; Meiji Seika, Tokyo, Japan). Then, they were housed individually and were allowed to recover for at least 2 days before habituation to treadmill running.

### 2.5 Habituation to treadmill running

All rats were habituated to running on a treadmill (KN-73, Natsume Seisakusho Co, Ltd., Tokyo, Japan) for a total of five sessions over 6 days. The running duration was 30 min/day, and the running speed was gradually increased from 5 to 25 m/min with no incline ^8,23^. Electrical shock grids were placed at the rear end of the treadmill and provided mild but aversive foot shocks (30 V) to rats to encourage running at the set treadmill speed. To minimize any stress derived from the shocks, electric shocks were limited and a gentle tail touch was given when rats did not start running again. After the habituation period, rats were able to run at the appropriate speeds even when the electric grid was turned off.

### 2.6 Treadmill running tests

On the test day, all rats fasted for 2 h before running to obtain stable metabolic conditions. In Experiments 1 and 2, treadmill running tests were conducted 2-3 days after the surgery for inserting the jugular catheter. In Experiment 1, an incremental exercise test was performed to assess the running speed at LT. Rats were forced to run to exhaustion; the running speed was started at 5 m/min and increased by 2.5 m/min every 3 minutes. The speed at which the LT was reached was determined according to the method by Beaver *et al..*^80^ In Experiment 2, a single 30-minute bout of running was conducted to verify the validation of exercise intensity based on the speed at LT as determined in Experiment 1. Rats were randomly subjected to run at 15 m/min (Light) or 25 m/min (Vigorous), or they were placed on a treadmill as the at-rest Control for 30 minutes. In both experiments, serial blood samples (0.1 ml) were taken at designated time points. Blood lactate and glucose were measured using an automated glucose–lactate analyzer (2300 Stat Plus, Yellow Springs Instruments, OH, USA).

In Experiment 3, rats were returned to their home cage after the test, rested for 90 min, and then were transcardially perfused with 0.9% saline, followed by 4% paraformaldehyde (PFA) in a 0.1 M phosphate buffer (PB; pH 7.4) under deep isoflurane anesthesia. The brains were sampled for immunohistochemical analysis. In Experiment 4, a microdialysis probe (2 mm membrane: FX-I-6-02, Eicom) was inserted into the dorsal hippocampus using a guide cannula 1 day before the test. The probe was connected to the microsyringe pump (ESP-32, Eicom) using a Teflon tubing (JT-10, Eicom) and perfused with Ringer’s solution (147 mM NaCl, 4 mM KCl, and 2.3 mM CaCl2) at 1.0 μl/min 2-3 h before the start of the test because a stable dialysate monoamine level is usually obtained 2 h after insertion of the probe. Using a fraction collector (EFC-82, Eicom), 15 μl of dialysate was collected every 15 minutes and 5 μl of 0.1M PB (pH 3.5) was added to ensure stable monoamine detection using HPLC with an electrochemical detector (HPLC-ECD). At the end of the experiments, rats were transcardially perfused with 0.9% saline, followed by 4% PFA in a 0.1 M PB under deep isoflurane anesthesia and the brain was collected to verify the position of the probe in coronal sections with DAPI (4’,6-diamidino-2-phenylindole). Please note that 7 rats in Experiment 4 were excluded from the analysis for the following reasons: failed insertion of the probe into the dorsal hippocampus (1 rat), detachment of the fixed probe during running (3 rats), and failed detection of peaks in chromatography (3 rats).

### 2.7 Immunohistochemistry

The brain was post-fixed for 24 h in 4% paraformaldehyde (PFA, 02890-45, Nacalai Tesque, Tokyo, Japan) in a 0.1 M PB and cryoprotected by immersion in a 0.1 M PB containing 20% sucrose for 1 day and 30% sucrose for 2-3 days. Frozen coronal brain sections of 30 µm thickness were cut with a microtome (REM-710, Yamato Kohki Industrial Co., Ltd., Saitama. Japan). Sections were washed with a 0.1M PB and blocked with a 0.1 M PB containing 1% Triton X-100 (35501-15, Nacalai Tesque) (PBT) plus 1% bovine serum albumin (01863-77, BSA; Nacalai Tesque). Then, slices were incubated with the designated primary antibodies in PBT-BSA for 24 h at 4 °C. The primary antibodies used in this study were rabbit polyclonal antibody against c-Fos (1:5,000; Cat# 226-003, RRID: AB_2231974, Synaptic Systems, GAU, Germany), mouse monoclonal antibody against NeuN (1:1,000; Cat# MAB377, RRID: AB_2298772, Millipore, MA, USA), mouse monoclonal antibody against tyrosine hydroxylase (1:2,000; Cat# F11:sc-25269, RRID: AB_628422, Santa Cruz, CA, USA), and goat polyclonal antibody against 5-HT (1:1,000; Cat# 20079, RRID: AB_572262, Immunostar Inc., WI, USA). After repeated washings with PBT-BSA, sections were incubated in a dark space with designated secondary antibodies diluted in PBT-BSA for >16 h at 4 °C. The secondary antibodies used in this study were Alexa Fluor 488-conjugated goat anti-rabbit IgG (1:5,000; Cat# A-11008, RRID: AB_143165, Invitrogen; Thermo Fisher Scientific Inc., MA, USA), Alexa Fluor 488-conjugated donkey anti-rabbit IgG (1:5,000; Cat# ab150073, RRID: AB_2636877, abcam, Cambridge, UK), Alexa Fluor 594-conjugated donkey anti-goat IgG (1:1,000; Cat# A-11058, RRID: AB_142540, Invitrogen; Thermo Fisher Scientific Inc., MA, USA), and AMCA AffiniPure donkey anti-mouse IgG (1:1,000; Cat# 715-155-150, RRID: AB_2340806, Jackson ImmunoResearch Laboratories, Inc., PA, USA). For DAPI staining to verify the location of microdialysis probes, slices were washed three times in a 0.1 M PB and then incubated with DAPI (1:1,000; 28718-90-3, Dojindo Laboratory, Kumamoto, Japan) in PBT-BSA for 10 minutes in the dark. Slices were washed three times in a 0.1 M PB, mounted on subbed slides, air dried, and coverslipped using a mounting medium.

### 2.8 Counts and quantification

All fluorescence z-stacked images were examined on an all-in-one microscope (BZ-X710, Keyence, Osaka, Japan) with a 20× objective lens. Manual counts of neurons were performed by an observer blinded to the group allocations. The number of c-Fos and NeuN double-positive neurons (c-Fos^+^/NeuN^+^) in the dorsal hippocampal dentate gyrus (DG), cornu ammonis (CA) 1, CA2, and CA3 sub-regions were counted hemilaterally in 4 coronal sections (from Bregma -3.60 mm, -4.16 mm, -5.20 mm, -5.80 mm). Total neuron numbers were normalized based on NeuN^+^ region volume (mm^3^), which was calculated by the area (mm^2^) multiplied by the thickness of sections (30 μm) using a BZ-X analyzer software (https://www.keyence.co.jp/products/microscope/fluorescence-microscope/bz-x700/models/bz-h3a/, version 1.4.1.1, RRID:SCR_017375). The numbers of c-Fos and TH or 5-HT double-positive neurons (c-Fos^+^/TH^+^ or c-Fos^+^/5-HT^+^) were bilaterally counted 120 μm apart throughout the LC (around Bregma -9.16 mm ∼ 10.52 mm, 10 ∼ 16 coronal sections) and 360 μm apart throughout the VTA, SNpc (around Bregma -4.80 mm ∼ -6.72 mm, 5 coronal sections), DRN, and MRN (around Bregma - 7.30 mm ∼ -8.80 mm, 5 coronal sections), all of which are known as monoaminergic nuclei projecting to the dorsal hippocampus.

### 2.9 HPLC-ECD for monoamine detection in the dorsal hippocampus

Dialysate samples collected from the probe at the dorsal hippocampus were injected into an HPLC-ECD system (HTEC-500; Eicom) using an autosampler (M-510; Eicom) and analyzed for NA, DA, and 5-HT concentration using the HPLC system with an EICOMPAK CAX column (2.0 mm, i.d. × 200 mm; Eicom) and a graft electrode (WE-3G; Eicom) set at 450 mV (vs Ag/AgCl reference electrode). The flow rate and injection volume were set to 250 μl/min and 17 μl, respectively. The mobile phase contained a 0.1 M ammonium acetate buffer, 0.05 mg/l sodium sulfate, 50 mg/l EDTA, and methanol (7:3, v/v), with a pH of 6.0. Data analysis was performed using Eicom EPC-710 software. Basal concentrations of the three monoamines (i.e., NA, DA, and 5-HT) were estimated from the averages at four HPLC time points before the beginning of running.

### 2.10 Statistical analyses

All data are expressed as mean ± standard error (SEM) and figures are generated using GraphPad Prism (https://www.graphpad.com/, version 8, RRID:SCR_002798, GraphPad Software, San Diego, CA, USA). All data were analyzed using R software (https://www.r-project.org/, version 4.2.3, RRID:SCR_001905). Assessment of the normality (Shapiro-Wilk test) of the data, equality of variance (Levene’s test), and sphericity (Mendoza’s test of sphericity) for repeated measures (RM) were performed. Statistical significance for parametric data was analyzed using one-way analysis of variance (ANOVA) followed by Shaffer’s *post-hoc* test, but Welch’s ANOVA was performed if equality of variance was not assumed. For the analysis of non-parametric data, the Kruskal-Wallis test with the Mann-Whitney U test adjusted by Holm’s post-hoc test was applied. For RM, two-way ANOVA followed by Shaffer’s post-hoc test was applied and the degree of freedom was adjusted using Huynh-Feldt-Lecoutre’s epsilon for non-spherical data. Pearson’s correlation coefficient was used for analyzing the strength of relationships. No tests for outliers were performed. Statistical significance was assumed at *p* < 0.05. No statistical methods were used to predetermine sample size, but the sample sizes were comparable to those reported in previous studies.^7,23,75–77^

## 3 Results

### 3.1 Measurement of LT and validation of exercise intensity

Initially, we developed a varied-intensity treadmill running model with adult male Long-Evans rats using LT as a signpost for exercise-induced stress. Our analysis revealed that the running speed at the LT of rats was 21.87 m/min ± 1.02 (*n* = 6; Fig. 1B, C). Thus, we defined 15 m/min, which is below the LT, as the speed for light-intensity running and 25 m/min, which is above the LT, as the speed for vigorous-intensity running. The observation of a significant increase in blood lactate concentration after a 30-minute session of vigorous-intensity running (two-way RM ANOVA, Shaffer’s *post hoc* test, group and time interaction, *F* _(2, 15)_ = 66.036, *p* < 0.001, Fig 1E) was sufficient to validate the varied-intensity running model based on LT.

**Figure 1.**
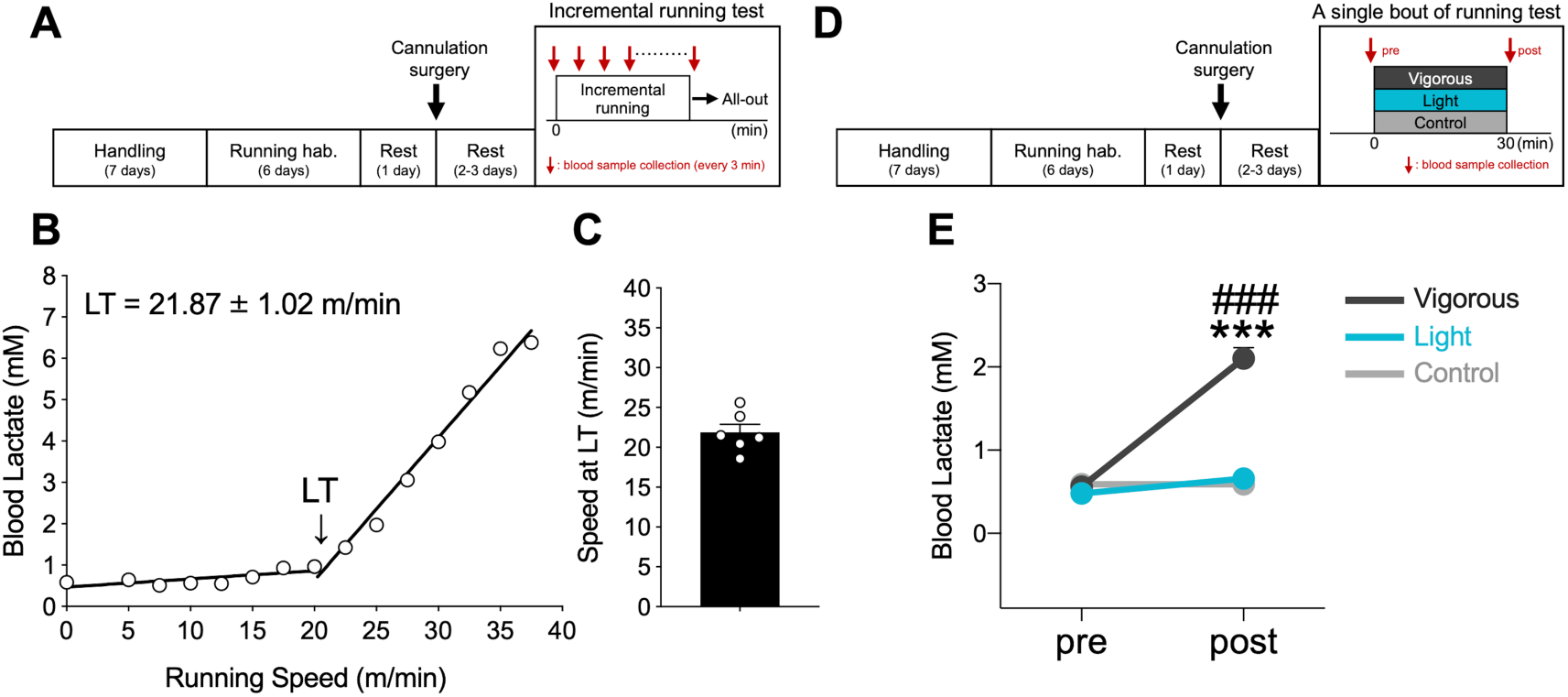
Measurement of lactate threshold and validation of exercise intensity. (A) Experimental design for incremental running test. (B) A representative LT profile from the incremental running test. (C) Speed of rats at LT was 21.87 ± 1.02 m/min (*n* = 6). (D) Experimental design for a single bout of the running test. We defined 15 m/min for light-intensity running and 25 m/min for vigorous-intensity running. (E) Blood lactate concentration was significantly increased only after a 30-minute session of vigorous-intensity running. All data are expressed as mean ± SEM. \*\*\**p* < 0.001 vs Control, ^###^*p* < 0.001 vs Light, *n* = 5-7/group. Control: sedentary control; Light: 15 m/min running; Vigorous: 25 m/min running; hab.: habituation.

### 3.2 Light-intensity-running-induced neuronal activation of dorsal hippocampus

We assessed dorsal hippocampal neuronal activation during different intensities of running using c-Fos immunostaining. Representative images in Figure 2B demonstrate that c-Fos expression in dorsal hippocampal sub-regions, including the DG, CA1, and CA3, significantly increased regardless of running speed. Conversely, the CA2 region was not affected by running (DG: Welch’s one-way ANOVA, Shaffer’s *post hoc* test, *F* _(2, 18)_ = 12.876, *p* < 0.001, Fig. 2C; CA1: one-way ANOVA, Shaffer’s *post hoc* test, *F* _(2, 18)_ = 9.994, *p* < 0.001, Fig. 2D; CA2: one-way ANOVA, Shaffer’s *post hoc* test, *F* _(2, 18)_ = 0.173, *p* = 0.843, Fig. 2E; CA3: one-way ANOVA, Shaffer’s *post hoc* test, *F* _(2, 18)_ = 11.256, *p* < 0.001, Fig. 2F).

**Figure 2.**
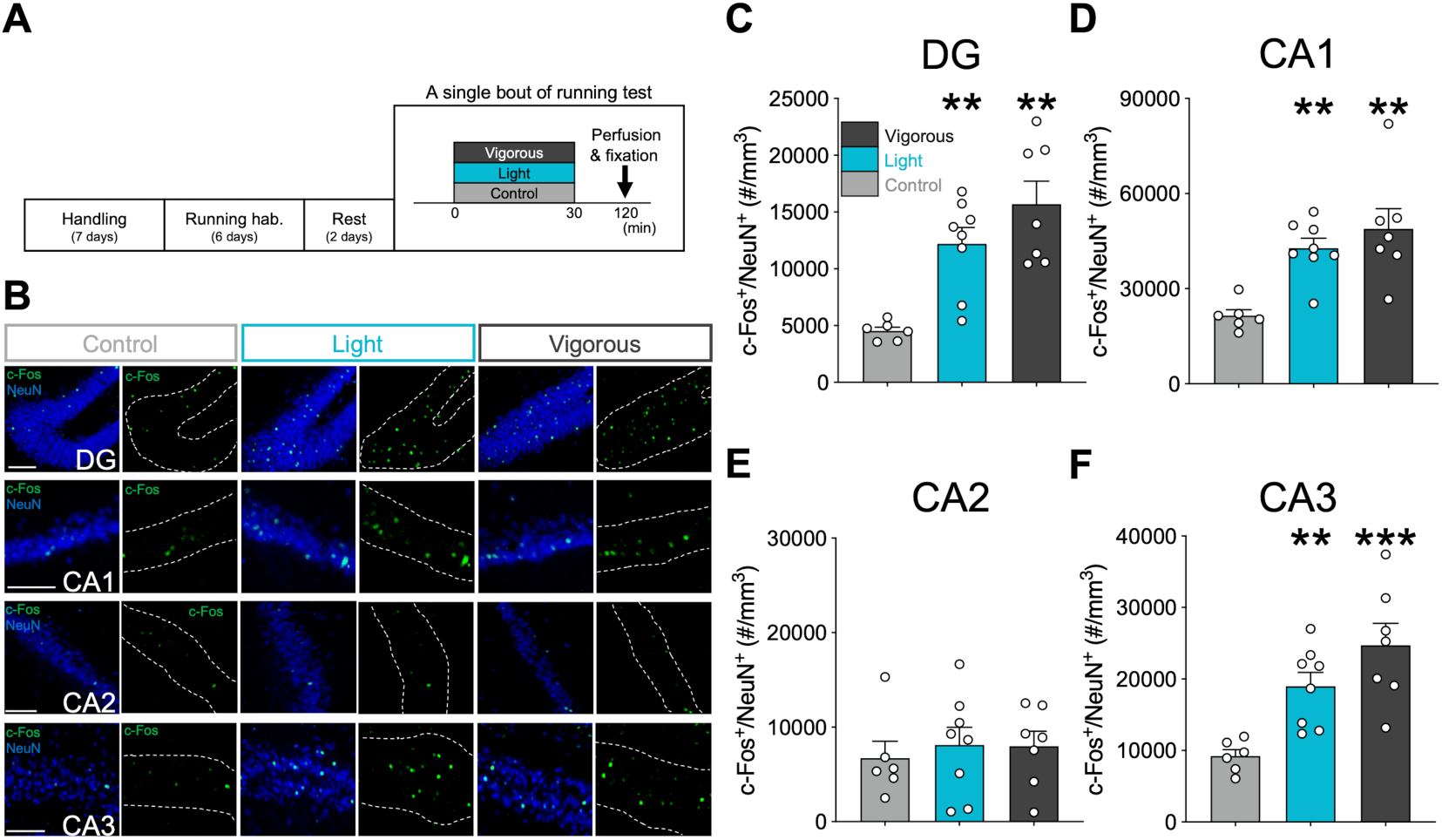
Running-induced neuronal activation at the dorsal hippocampus during varied-intensity treadmill running. (A) Experimental design for a single bout of the running test for immunohistochemical analysis. (B) Representative double-fluorescence images of c-Fos (green) and NeuN (blue) in the DG, CA1, and CA3 regions of the dorsal hippocampus. Scale bars show 100 μm. c-Fos expressions in the DG (C), CA1 (D), CA2 (E), and CA3(F) were significantly increased with running exercise regardless of running speed (light or vigorous). All data are expressed as mean ± SEM. \*\**p* < 0.01, \*\*\**p* < 0.001 vs Control, *n* = 6-8 / group. Control: sedentary control; Light: 15 m/min running; Vigorous: 25 m/min running; hab.: habituation.

### 3.3 Light-intensity running increased dorsal hippocampal NA and DA release

To assess the monoaminergic neurotransmission to the dorsal hippocampus, we measured extracellular monoamine levels in the dorsal hippocampus using *in vivo* microdialysis and HPLC methods. We found that treadmill running induced a significant increase of both NA and DA levels in the dorsal hippocampus depending on running speed (NA: two-way RM ANOVA, Shaffer’s *post hoc* test, group, *F* _(2, 17)_ = 11.545, *p* < 0.001, time, *F* _(3.5, 59.58)_ = 16.236, *p* < 0.001, group and time interaction, *F* _(7.01, 59.58)_ = 5.393, *p* < 0.001; DA: two-way RM ANOVA, Shaffer’s *post hoc* test, group, *F* _(2, 17)_ = 18.720, *p* < 0.001, time, *F* _(4.01, 68.23)_ = 14.019, *p* < 0.001, group and time interaction, *F* _(8.03, 68.23)_ = 4.959, *p* < 0.001, Fig. 3E, F). Notably, DA release was significantly sustained into the post-running time period. Dorsal hippocampal 5-HT levels, on the other hand, were not affected by running (two-way RM ANOVA, Shaffer’s *post hoc* test, group, *F* _(2, 17)_ = 0.046, *p* = 0.955, time, *F* _(2.79, 47.46)_ = 16.236, *p* < 0.05, group and time interaction, *F* _(5.58, 47.46)_ = 1.16, *p* = 0.343, Fig. 3G).

**Figure 3.**
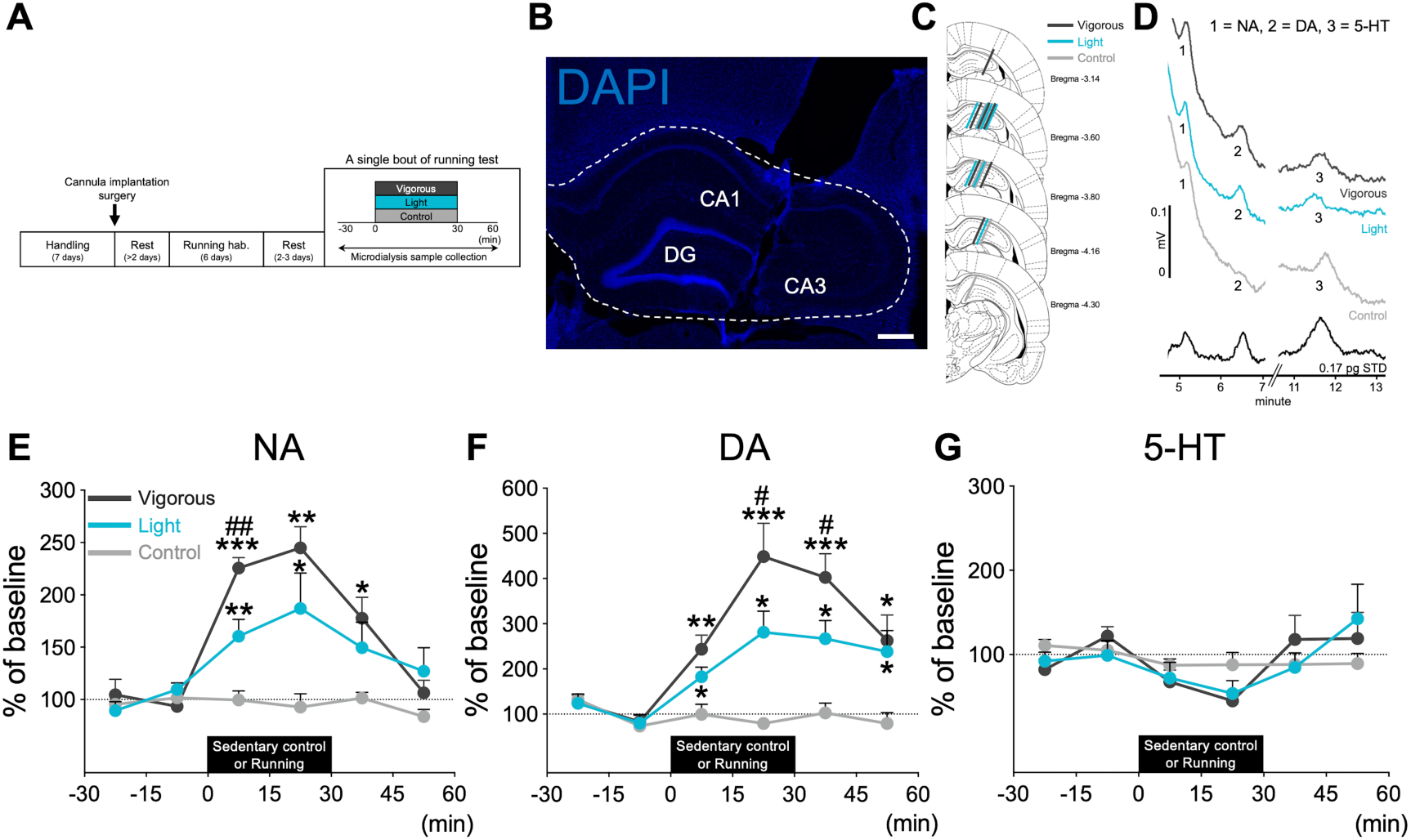
Running increases dorsal hippocampal NA and DA release, but not 5-HT release. (A) Experimental design for a single bout of the running test for dorsal hippocampal monoamine detection with *in vivo* microdialysis. (B) Representative DAPI fluorescence image of microdialysis probe position in the dorsal hippocampus. Scale bars show 500 μm. (C) Schematic representations indicating the location of the microdialysis probe. Each bar represents the probe position of an individual rat, and the color of the bar corresponds to the experimental group (Control: light gray, Light: blue, and Vigorous: dark gray). (D) Representative chromatograms showing NA (peak 1), DA (peak 2), and 5-HT (peak 3) from Control (light gray), Light (blue), and Vigorous (dark gray). Treadmill running induced a significant increase of both NA (E) and DA (F) levels in the dorsal hippocampus, depending on running speed, but not 5-HT (G). All data are expressed as mean ± SEM. \**p* < 0.05, \*\**p* < 0.01, \*\*\**p* < 0.001 vs Sed, ^#^*p* < 0.05, ^##^*p* < 0.01 vs Light, *n* = 6-7/group. Control: sedentary control; Light: 15 m/min running; Vigorous: 25 m/min running; hab.: habituation.

### 3.4 Running stimulated TH^+^ neurons in LC and VTA, but not raphe 5-HT^+^ neurons

To examine the responsiveness of brainstem monoaminergic nuclei, we performed double-immunostaining of c-Fos^+^/TH^+^ neurons for the activation of NA- and DA-producing neurons, and c-Fos^+^/5-HT^+^ neurons for the activation of serotonergic neurons, as shown in Figure 4A-C. TH^+^ neurons in the LC (LC^TH^) and VTA (VTA^TH^), which are known as the main sources of dorsal hippocampal NA and DA, respectively, were significantly stimulated by both light- and vigorous-intensity running (LC: one-way ANOVA, Shaffer’s *post hoc* test, *F* _(2, 18)_ = 20.908, *p* < 0.001, Fig. 4D; VTA: one-way ANOVA, Shaffer’s *post hoc* test, *F* _(2, 18)_ = 4.992, *p* < 0.05, Fig. 4E). However, running did not have any impact on dopaminergic neurons in the SNpc (SNpc^TH^ neurons), or on serotonergic neurons in the DRN and MRN (DRN^5-HT^ neurons and MRN^5-HT^ neurons, respectively) (SNpc: Kruskal-Wallis test, Mann-Whitney U test adjusted by Holm’s *post-hoc* test, *X*^2^ = 4.653, df = 2, *p* = 0.097, Fig. 4F; DRN: one-way ANOVA, Shaffer’s *post hoc* test, *F* _(2, 18)_ = 0.832, *p* = 0.454, Fig. 4G; MRN: one-way ANOVA, Shaffer’s *post hoc* test, *F* _(2, 18)_ = 1.640, *p* = 0.222, Fig. 4H). Note that TH^+^ neurons in the retrorubral field (RRF) is also reported to have projections to the hippocampus,^81^ which were not activated by running (Fig. S2B). Additionally, TH^+^ neurons were also observed in the rostral region of the DRN (DRN^TH^ neurons), which are reported to contribute to wakefulness by salience stimuli, opioid addiction, and pain modulation through the release of DA.^82^ In this study, however, running did not affect the c-Fos expression in DRN^TH^ neurons (Fig. S2D).

**Figure 4.**
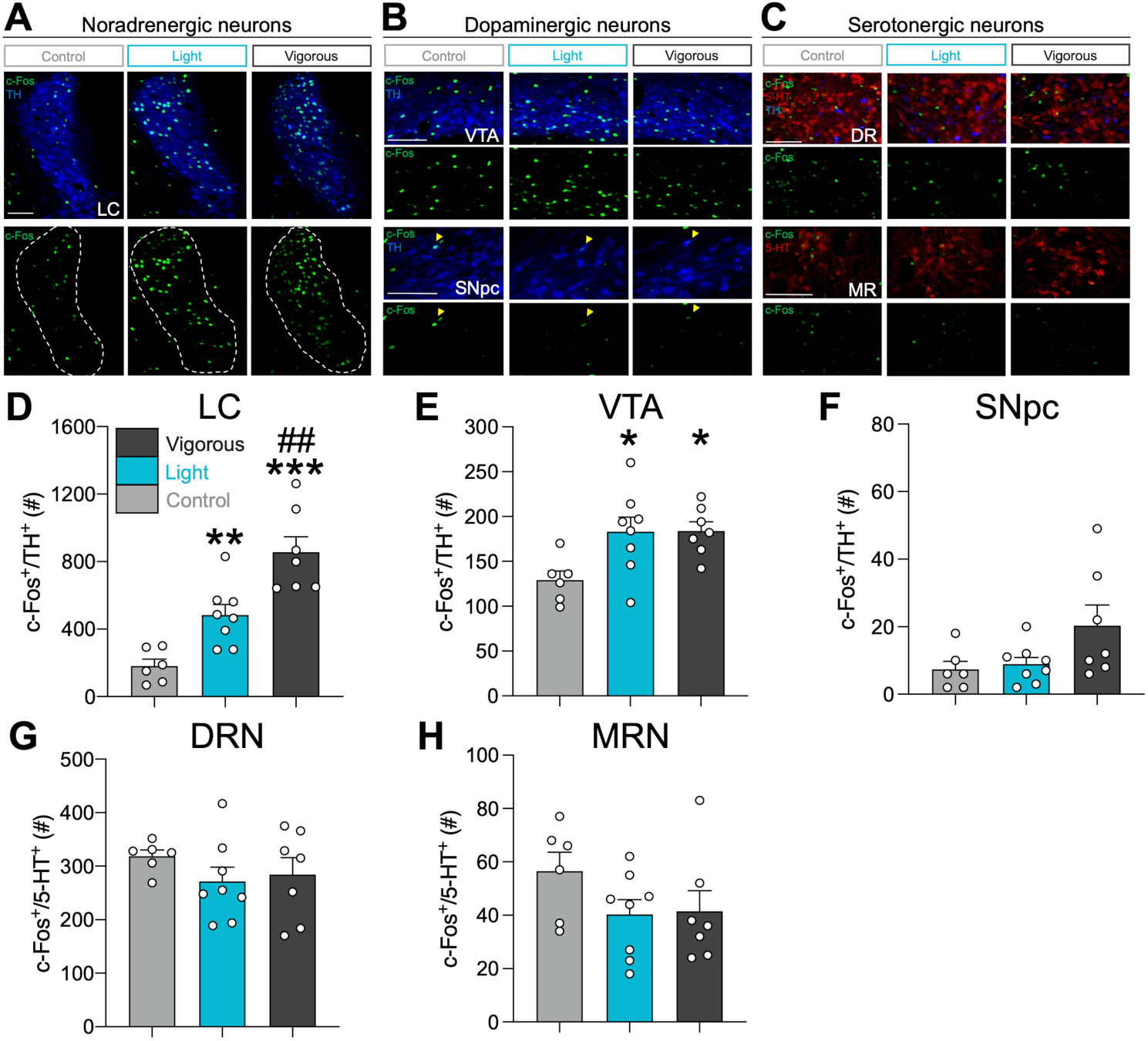
Running stimulates LCTH and VTATH neurons, but not raphe 5-HT+ neurons. Representative double-fluorescence images of c-Fos (green), TH (blue), and 5-HT (red) neurons in LC (A); VTA and SNpc (B); and DRN and MRN (C). Scale bars show 100 μm. Yellow heads show c-Fos^+^/TH^+^ neurons. LC^TH^ and VTA^TH^ neurons were significantly stimulated in both running groups (D, E). However, running did not have an impact on SNpc^TH^, DRN^5-HT^, and MRN^5-HT^ neurons (F, G, H). All data are expressed as mean ± SEM. \**p* < 0.05, \*\**p* < 0.01, \*\*\**p* < 0.001 vs Control, ^##^*p* < 0.01 vs Light, *n* = 6-8 / group. Control: sedentary control; Light: 15 m/min running; Vigorous: 25 m/min running.

### 3.5 Neuronal associations between dorsal hippocampus and brainstem monoaminergic nuclei

To explore the potential association between light-exercise-induced neuronal activities in brainstem monoaminergic nuclei and dorsal hippocampal activation, we conducted Pearson’s correlation analysis using the dataset from the control group and the light-intensity running group. This analysis assessed the correlation between c-Fos expression in the brainstem monoaminergic nuclei and the dorsal hippocampus. A correlation matrix heatmap that summarizes the correlation results is shown in Figure 5. c-Fos expression in LC^TH^ neurons exhibited a significant and positive correlation within all examined dorsal hippocampal sub-regions except CA2 (*n* = 14, DG: *r* = 0.607, *p* < 0.05, Fig 5B; CA1: *r* = 0.691, *p* < 0.01, Fig 5C; CA2: *r* = 0.132, *p* = 0.652; CA3: *r* = 0.643, *p* < 0.05, Fig 5D).

**Figure 5.**
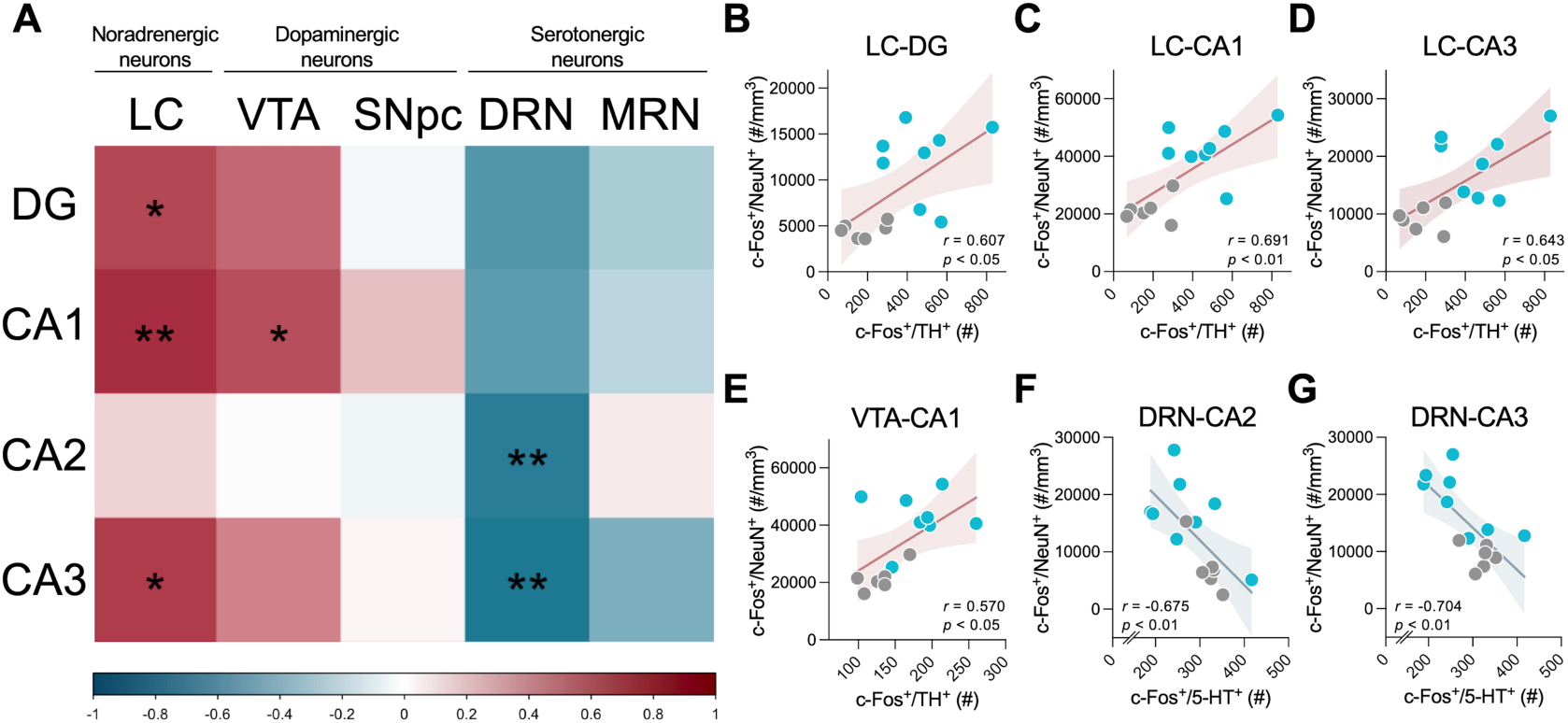
Neuronal associations between dorsal hippocampus and brainstem monoaminergic nuclei. (A) A correlation matrix heatmap is shown to summarize the correlation analysis between c-Fos expression in dorsal-hippocampal sub-regions and that in LC^TH^, VTA^TH^, SNpc^TH^, DRN^5-HT^, and MRN^5-HT^ neurons. Colors in the scale show correlation strength. c-Fos expression in LC^TH^ neurons exhibited a significant and positive correlation within the dorsal DG (B), CA1 (C), and CA3 (D). A significant positive correlation was also found between c-Fos expression in VTA^TH^ neurons and in the dorsal CA1 (E). c-Fos expression in DRN^5-HT^ neurons was significantly and negatively correlated with that in CA2 (F) and CA3 (G). The colored line represents linear regression, and the colored band represents 95% confidence bands. \**p* < 0.05, \*\**p* < 0.01. *n* = 14.

Additionally, a significant positive correlation was found between c-Fos expression in VTA^TH^ neurons and in the dorsal CA1 (VTA: *n* = 14, DG: *r* = 0.482, *p* = 0.0816; CA1: *r* = 0.570, *p* < 0.05, Fig 5E; CA2: *r* = 0.004, *p* = 0.989; CA3: *r* = 0.391, *p* = 0.167). The higher level of c-Fos expression in DRN^5-HT^ neurons was significantly correlated with lower levels of CA2 and CA3 c-Fos expression (*n* = 14, DG: *r* = -0.532, *p* = 0.0502; CA1: *r* = -0.506, *p* = 0.0649; CA2: *r* = -0.675, *p* < 0.01, Fig 5F; CA3: *r* = -0.704, *p* < 0.01, Fig 5G). However, no correlation was found in the c-Fos expression between SNpc^TH^ or MRN^5-HT^ neurons and dorsal hippocampal sub-regions (SNpc: *n* = 14, DG: *r* = -0.0362, *p* = 0.902; CA1: *r* = 0.196, *p* = 0.502; CA2: *r* = -0.0477, *p* = 0.871; CA3: *r* = 0.0374, *p* = 0.899; MRN: *n* = 14, DG: *r* = -0.252, *p* = 0.385; CA1: *r* = -0.207, *p* = 0.478; CA2: *r* = 0.0693, *p* = 0.814; CA3: *r* = -0.402, *p* = 0.155).

## 4. Discussion

To probe the neural circuitry responsible for activating the dorsal hippocampus during light exercise and its subsequent contribution to enhanced memory functions, this study examined the involvement of the ascending monoaminergic system—a critical neuromodulatory mechanism for memory—in the activation of the dorsal hippocampus induced by light exercise. Employing a rat exercise model based on the LT, we assessed neuronal activity in the dorsal hippocampus and brainstem monoaminergic neurons and observed real-time dynamics of monoamine release in the dorsal hippocampus using *in vivo* microdialysis. The results demonstrate that even light exercise led to increased c-Fos expression in specific subregions of the dorsal hippocampus and triggered the release of both NA and DA within this brain region. Furthermore, the activity of LC^TH^ neurons and VTA^TH^ neurons elevated during exercise even at the light-intensity, and both were positively correlated with the activation of the dorsal hippocampus. In contrast, the serotonergic system did not appear to be influenced by exercise. These findings imply LC-noradrenergic and VTA-dopaminergic contributions to dorsal hippocampal activation during light exercise (Fig. 6), offering insights into the circuitry behind light-exercise-enhanced learning and memory.

**Figure 6.**
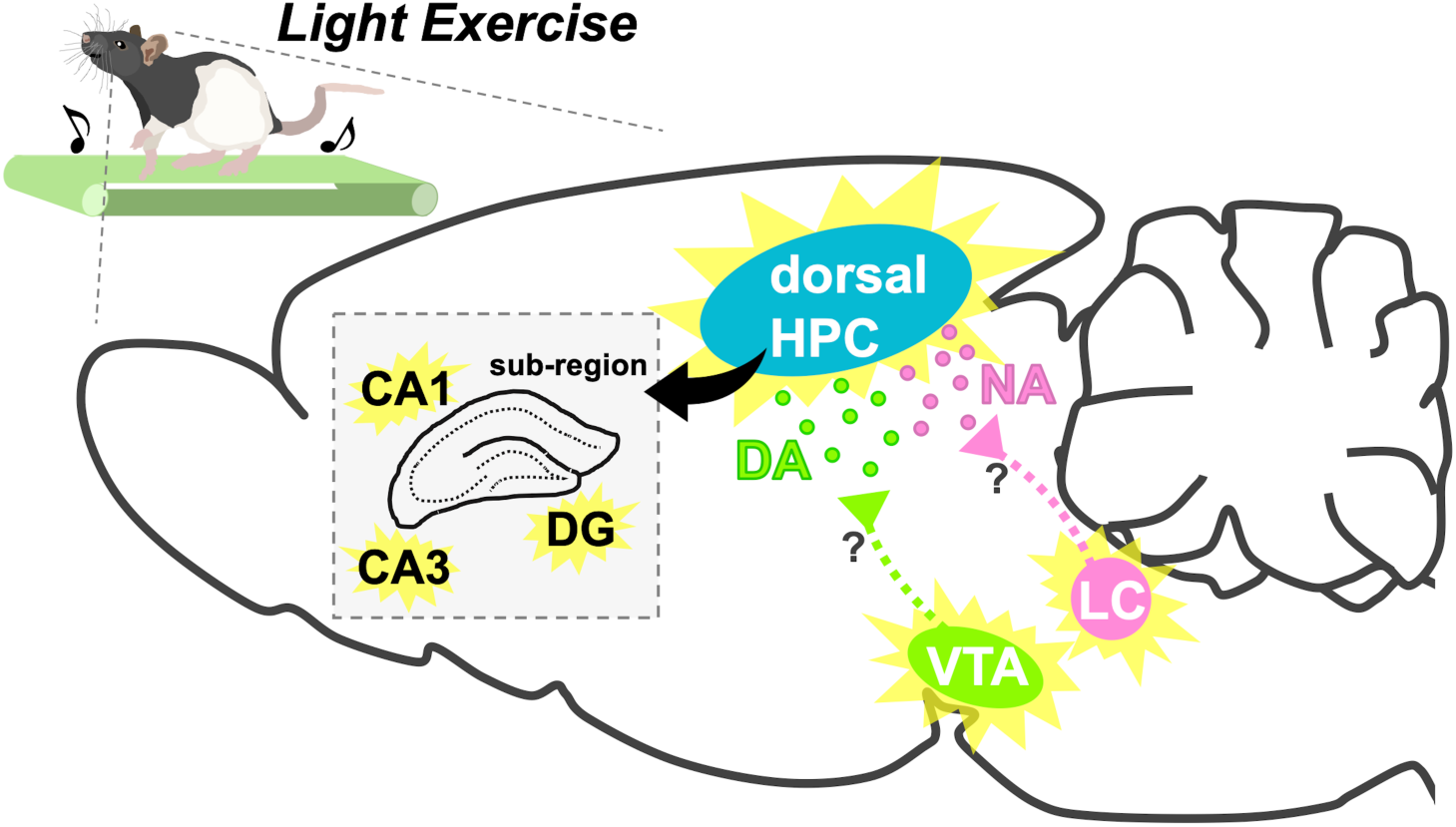
Schematic diagram showing possible circuit mechanism for light-exercise-induced dorsal hippocampal activation. Light exercise activates TH^+^ neurons at the LC and the VTA. At the dorsal hippocampus, subsequently, axon terminals originating from the LC^TH^ and the VTA^TH^ increase the release of NA and DA, respectively. These, in turn, upregulate the neuronal activity of the dorsal hippocampal DG, CA1, and CA3 sub-regions. HPC, hippocampus; DG, dentate gyrus; NA, noradrenaline; DA, dopamine; LC, locus coeruleus; VTA, ventral tegmental area.

Here, we first established a varied-intensity treadmill running model for male adult Long-Evans rats based on LT as a physiological index of running intensity and stress. The speed at the LT was similar to that in our previous work using male adult Wistar rats (Long-Evans: 21.87 m/min ± 1.02 vs Wistar: 20.7 m/min ± 2.1; Fig. 1B, C).^23^ In addition, a significant increase in blood lactate concentration was observed following a single bout of vigorous-intensity running, but not with light-intensity running (Fig. 1E), which is consistent with our previous reports.^23,25^ These results confirm the validity of our experimental model (LT-based varied-intensity running) and demonstrate that the two running groups exhibited different metabolic and stress-related responses.

Previous studies have reported that acute varied-intensity treadmill running activates the dorsal hippocampus, including the DG, CA1, and CA3 sub-regions, even at low-speeds.^6,7,9^ Our current results support these findings since both light- and vigorous-intensity running increased c-Fos expression in the dorsal DG, CA1, and CA3 sub-regions (Fig. 2C, D, F). In contrast, the dorsal CA2 region was not activated with running (Fig. 2E). The CA2 is known as a small hippocampal sub-region that modulates social memory and that is stimulated by novel encounters.^83^ Since all rats were well familiarized with the experimental conditions and there were no novel or social stimuli, it is reasonable that activity in the CA2 region was not affected. Additionally, we examined the activation of the ventral hippocampus following running and found an increase of c-Fos expression across all sub-regions of the ventral hippocampus in both running groups (Fig. S1). These results suggest that light-intensity running is sufficient to elevate the neuronal activity in the hippocampus across the dorsal-to-ventral axis, which may help to explain the positive effects of exercise on dorsal-hippocampus-related functions, such as memory, as well as ventral-hippocampus-related anxiolytic and antidepressant effects.^3,29,84^

Our *in vivo* microdialysis experiment provides the first evidence that even light-intensity running increases the release of NA and DA at the dorsal hippocampus (Fig. 3E, F). Given that NA and DA both perform critical roles in dorsal hippocampal neuronal activity, plasticity, and memory process,^60,61,63,64,85–87^ our results suggest that the dorsal hippocampal NA and DA release contributes to the light-exercise-induced hippocampal activation and beneficial effects, such as hippocampal neurogenesis and spatial memory performance.^9,29,30^ Notably, NA and DA have been recognized as important factors in memory consolidation,^63,86^ as shown by the synaptic tagging and capture hypothesis,^88,89^ raising a hypothesis that a single-bout of light-intensity running could facilitate memory retention through noradrenergic and dopaminergic dorsal hippocampal activation. Intriguingly, prior studies demonstrate that a single session of vigorous-intensity exercise elevates NA and DA concentrations in the hippocampus and promotes novel object recognition, which is a dorsal-hippocampus-related function, through the β adrenergic receptor and dopamine D1 receptor, respectively.^3,79^ Although we did not directly examine this in the present study, these findings support our hypothesis.

We confirmed that treadmill running, even at light intensity, activates LC^TH^ and VTA^TH^ neurons (Fig. 4D, E). Furthermore, our correlation analysis observed that LC^TH^ activation positively correlates with dorsal hippocampal activation, and the activities of VTA^TH^ were positively associated with those of the dorsal CA1 region (Fig. 5). These results suggest that running stimulates the dorsal hippocampal neurons through noradrenergic projection from the LC and dopaminergic projection from the VTA. While our study didn’t investigate the projections from light-exercise-activated TH^+^ neurons in the LC and the VTA to the dorsal hippocampus, interestingly, a recent study with *in vivo* two-photon calcium imaging provides robust support for our hypothesis, which demonstrates an elevation in the activities of both the LC and the VTA axons within the dorsal CA1 in head-fixed mice during voluntary running.^90^ Both nuclei are known as major sources of dorsal hippocampal NA and DA,^64,65,91^ but recent studies have shown that LC terminals co-release DA and NA to the dorsal hippocampus, and that LC-DA, but not LC-NA, plays a significant role in spatial memory and learning of novel context.^63–66,85,92^ On the other hand, VTA^TH^ neurons have sparse input to the dorsal hippocampus and do not have a significant impact on spatial memory.^63,64^ However, hippocampal DA from VTA^TH^ neurons is still reported to be involved in hippocampus-related functions, such as spatial memory and aversive memory.^70,71,93^ The significance of LC^TH^ and VTA^TH^ neurons for the dorsal hippocampal functions remains elusive; thus, future studies using viral-tracing techniques and chemogenetic/optogenetic manipulation are needed to investigate the noradrenergic and dopaminergic mechanisms underlying the beneficial effects of light exercise.

Unlike the noradrenergic and the dopaminergic systems, 5-HT release was not increased at the dorsal hippocampus, nor were DRN^5-HT^ or MRN^5-HT^ neurons found to be stimulated by treadmill running (Fig. 3G; Fig. 4G, H). However, we found a negative correlation between c-Fos expression in the DG, CA2, and CA3 sub-regions and that in DRN^5-HT^ neurons (Fig. 5F, G). Previous studies have shown that running increases hippocampal neuronal activity and also facilitates hippocampal LTP induction.^94–96^ Conversely, the hippocampal 5-HT can exert inhibitory effects on basal neural activity and LTP induction.^97–99^ Considering these findings, our results imply that running enhances hippocampal neuronal activity and LTP induction through the inhibitory control of the serotonergic inhibitory projection from DRN^5-HT^ neurons to the dorsal hippocampus.

From a clinical perspective, light exercise, such as yoga, tai-chi, and slow running, is a simple and feasible intervention accessible by most people, including patients with mild cognitive impairment (MCI) and Alzheimer’s disease (AD). Recent studies have shown that the hippocampal NA and DA loss and degeneration of LC^TH^ and VTA^TH^ neurons and axons are found in AD model rodents and AD patients.^100–104^ Notably, selective accumulation of amyloid β and tau proteins are observed in LC neurons before the onset or at the early stage of AD symptoms.^103–107^ Light exercise has been reported to have a positive effect on MCI and AD.^108–110^ Our findings using a translational animal exercise model, therefore, could provide insight into understanding the mechanisms underlying the preventive or therapeutic role of light exercise in cognitive dysfunction conditions, such as AD.

## 5. Conclusion

Collectively, our findings using a translational exercise model revealed that even light-intensity running can stimulate the dorsal hippocampal neuronal activity and the release of NA and DA probably from LC and VTA, respectively but that it does not affect the serotonergic system. These findings suggest a possible contribution of LC-noradrenergic and VTA-dopaminergic projections to dorsal hippocampal activation during light exercise (Fig. 6), which may provide insight into the circuit mechanism that underlies light-exercise-enhanced learning and memory.

## Abbreviations

5-HT: serotonin
AAS: ascending arousal system
ACTH: adrenocorticotropic hormone
AD: Alzheimer’s disease
BDNF: brain-derived neurotrophic factor
BSA: bovine serum albumin
CA: cornu ammonis DA: dopamine
DAPI: 4’,6-diamidino-2-phenylindole DG: dentate gyrus
DRN: dorsal raphe nucleus
HPLC-ECD: high-performance liquid chromatography with an electrochemical detector
IGF-I: insulin-like growth factor I
LC: locus coeruleus
LTD: long-term depression
LTP: long-term potentiation
LT: lactate threshold
MCI: mild cognitive impairment
MRN: median raphe nucleus NA: noradrenaline
PB: phosphate buffer
PBT: phosphate buffer with Triton X-100
PFA: paraformaldehyde
PRP: plasticity-related protein
PVN: paraventricular nucleus
RM: repeated measures
RRF: retrorubral field
SNpc: substantia nigra pars compacta
TH: tyrosine hydroxylase
VTA: ventral tegmental area

## Author Contributions

Taichi Hiraga, Toshiaki Hata, Shingo Soya, Kanako Takahashi, Masahiro Okamoto, and Hideaki Soya conceived and designed the experiments; Joshua P. Johansen provided the experimental animals; Taichi Hiraga and Kanako Takahashi performed the treadmill running experiments for LT measurement and validation of exercise intensity; Taichi Hiraga performed the treadmill running experiments for immunohistochemistry and microdialysis; Toshiaki Hata and Ryo Shimoda performed experimental assistances; Taichi Hiraga and Ryo Shimoda analyzed the data; Taichi Hiraga wrote the original manuscript, and all authors were involved in reviewing and editing the manuscript.

## Acknowledgments

The authors are grateful to colleagues at the Laboratory of Exercise Biochemistry and Neuroendocrinology (Faculty of Health and Sport Sciences, University of Tsukuba, Japan), particularly R. Kuwamizu, H. Koizumi, T. Ferenc, F. Grenier, and M.J. Ikemoto for insightful scientific discussions. We thank J.P. Johansen (RIKEN CBS) for providing the experimental animals (TH-Cre Long-Evans rats line) and discussion. We would like to express our gratitude to M. Noguchi (ELCS English Language Consultation, Japan) for helping with the manuscript. This research was supported in part by KAKENHI

Grants-in-Aid for Scientific Research (A) (18H04081; 21H04858) (to H.S.); KAKENHI Grants-in-Aid for Scientific Research on Innovative Areas: Next Generation Exercise Program for Developing Motivation, Body and Mind Performance (16H06405) (to H.S.); Japan Science and Technology Agency (JST)-Mirai Program (JPMJMI19D5) (to H.S.); Grant-in-Aid for Japan Society for the Promotion of Science Fellowships (20J20887) (to T.H.).

## Supplemental data

Light-exercise-induced dopaminergic and noradrenergic stimulation in the dorsal hippocampus: using a rat physiological exercise model

Taichi Hiraga, Toshiaki Hata, Shingo Soya, Ryo Shimoda, Kanako Takahashi, Mariko Soya, Koshiro Inoue, Joshua P. Johansen, Masahiro Okamoto, and Hideaki Soya

Corresponding author: Hideaki Soya, Ph.D.

This PDF file includes:

- Figure S1 Running induces neuronal activation of the ventral hippocampus
- Figure S2 RRF^TH^ neurons and DRN^TH^ neurons are not activated by running.

---

**Figure S1.**
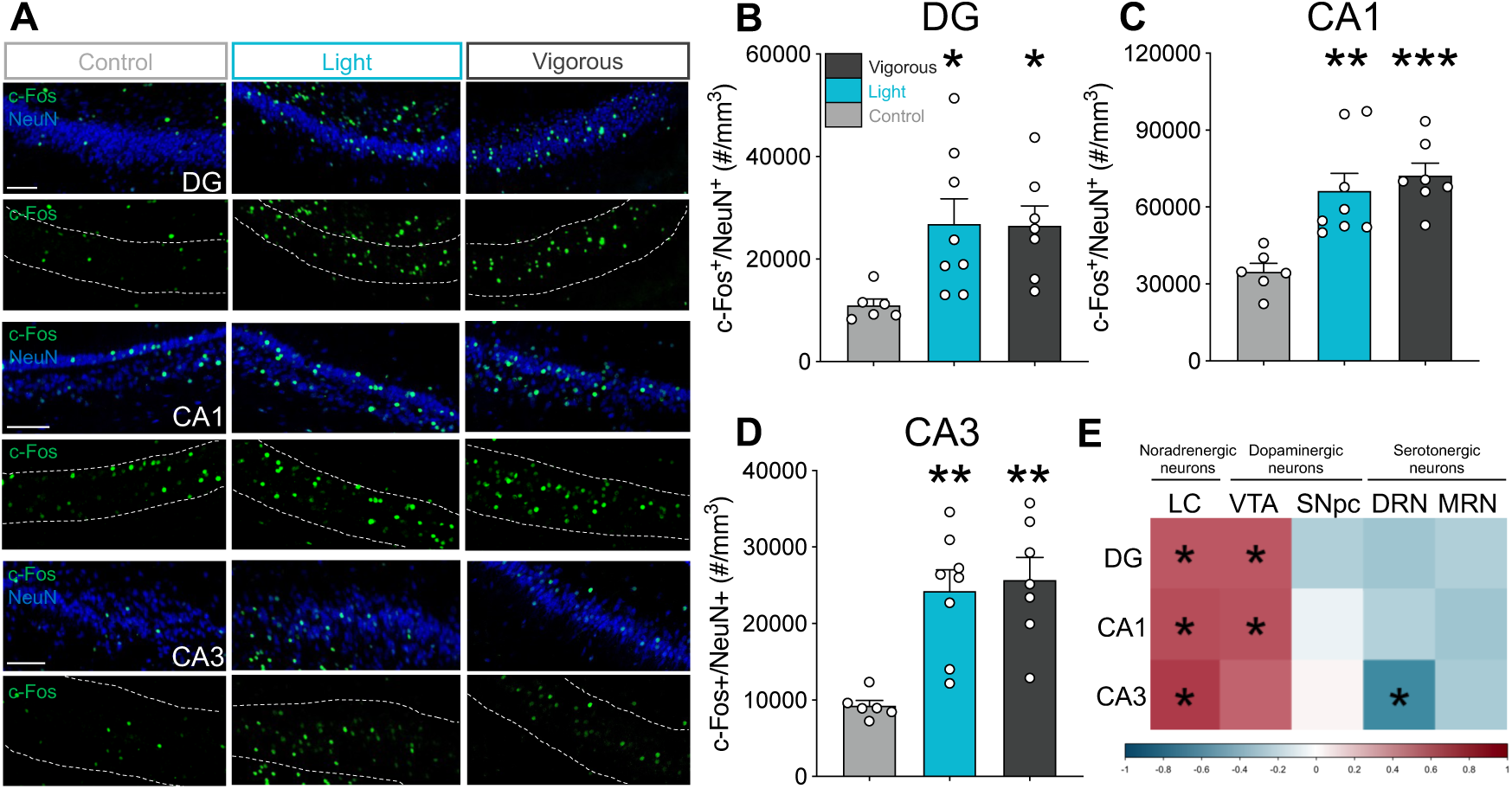
Running induces neuronal activation of the ventral hippocampus. (A) Representative double-fluorescence images of NeuN (blue) and c-Fos (green) in the DG, CA1, and CA3. Scale bars show 100 μm. c-Fos expression in ventral hippocampal sub-regions, including the DG (B), CA1 (C), and CA3 (D), was also significantly increased regardless of running speed (DG: Kruskal-Wallis test, Mann-Whitney U test adjusted by Holm’s *post-hoc* test, *X*^2^ = 10.245, df = 2, *p* < 0.01; CA1: one-way ANOVA, Shaffer’s *post hoc* test, *F* _(2, 18)_ = 11.443, *p* < 0.001; CA3: Welch’s one-way ANOVA, Shaffer’s *post hoc* test, *F* _(2, 18)_ = 11.758, *p* < 0.001). (E) A correlation matrix heatmap is shown to summarize the correlation results between c-Fos expression in monoaminergic nuclei and that in ventral hippocampal sub-regions. Colors in the scale show correlation strength. Data were analyzed using Pearson’s correlation. c-Fos expression in the LC^TH^ had a significant positive correlation with that in all ventral hippocampal sub-regions (*n* = 14, DG: *r* = 0.543, *p* < 0.05; CA1: *r* = 0.587, *p* < 0.05; CA3: *r* = 0.654, *p* < 0.05). Additionally, a significant positive correlation was found between c-Fos expression in the VTA^TH^ neurons and in the ventral DG and CA1 (*n* = 14, DG: *r* = 0.550, *p* < 0.05; CA1: *r* = 0.564, *p* < 0.01; CA3: *r* = 0.507, *p* = 0.064). The higher level of c-Fos expression in the DRN^5-HT^ neurons was significantly correlated with the lower level of ventral CA3 c-Fos expression (*n* = 14; DG: *r* = -0.085, *p* = 0.715; CA1: *r* = -0.090, *p* = 0.697; CA3: *r* = - 0.522, *p* < 0.05). No correlation was found between c-Fos expression in SNpc^TH^ or MRN^5-HT^ neurons and in ventral hippocampal sub-regions (SNpc: *n* = 14, DG: *r* = -0.234, *p* = 0.422; CA1: *r* = -0.051, *p* = 0.864; CA3: *r* = 0.039, *p* = 0.895; MRN: *n* = 14, DG: *r* = - 0.238, *p* = 0.412; CA1: *r* = -0.283, *p* = 0.328; CA3: *r* = -0.253, *p* = 0.382). All data are expressed as mean ± SEM. \**p* < 0.05, \*\**p* < 0.01, \*\*\**p* < 0.001 vs Control; *n* = 6-8/group. Control: sedentary control; Light: 15 m/min running; Vigorous: 25 m/min running.

**Figure S2.**
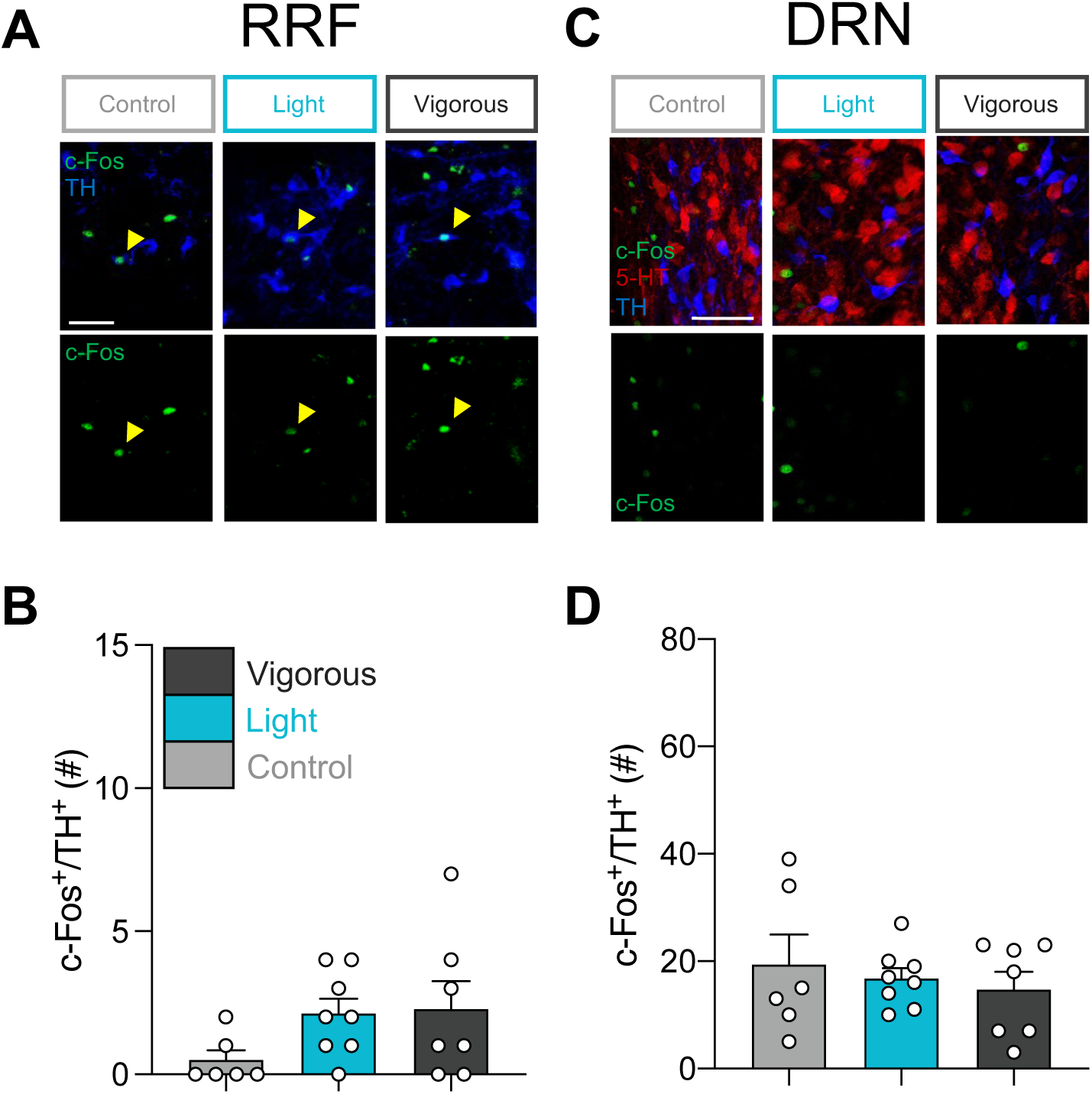
RRFTH neurons and DRNTH neurons are not activated by running. (A) Representative double-fluorescence images of c-Fos (green) and TH (blue) neurons in the RRF. Scale bars show 100 μm. Yellow heads show c-Fos^+^/TH^+^ neurons. (B) RRF^TH^ neurons were not activated by running (Kruskal-Wallis test, Mann-Whitney U test adjusted by Holm’s *post-hoc* test, *X*^2^ = 4.494, df = 2, *p* = 0.106). (C) Representative triple-fluorescence images of c-Fos (green), TH (blue), and 5-HT (red) neurons in the DRN. Scale bars show 50 μm. (D) DRN^TH^ neurons were not activated by running (Welch’s one-way ANOVA, Shaffer’s *post hoc* test, *F* _(2, 18)_ = 0.384, *p* =0.687). All data are expressed as mean ± SEM. *n* = 6-8/group. Control: sedentary control; Light: 15 m/min running; Vigorous: 25 m/min running.

